# Complete genome sequences of the species type strains *Sinorhizobium garamanticum* LMG 24692^T^ and *Sinorhizobium numidicum* LMG 27395^T^ and CIP 109850^T^

**DOI:** 10.1101/2023.04.25.538266

**Authors:** Sabhjeet Kaur, Daniel Espinosa-Sáiz, Encarna Velázquez, Esther Menéndez, George C diCenzo

**Author notes:** **Corresponding authors:** Esther Menéndez and George Colin diCenzo.

## Abstract

The genus *Sinorhizobium* comprises rhizobia that fix nitrogen in symbiosis with legumes. To support taxonomic studies of this genus and of rhizobia more broadly, we report complete genome sequences and annotations for the species type strains *Sinorhizobium garamanticum* LMG 24692^T^ and *Sinorhizobium numidicum* LMG 27395^T^ and CIP 109850^T^. Average nucleotide identity and core-genome phylogenetic analyses confirm that *S. garamanticum* and *S. numidicum* represent distinct species.

## Main text

The genus *Sinorhizobium* currently consists of 17 validly published species (1), all of which can enter into nitrogen-fixing symbiosis with leguminous plants. Prior to this work, the genome sequences of the species type strains for 14 of the 17 *Sinorhizobium* spp. were publicly available (1–9). Genomic data for species type strains is valuable in bacterial systematics by facilitating the delineation of species and higher-order groups (10). To support taxonomic studies of the genus *Sinorhizobium* and of rhizobia more broadly, we report the genome sequences of type strains for two additional *Sinorhizobium* species: *S. garamanticum* LMG 24692^T^ and *S. numidicum* LMG 27395^T^ and CIP 109850^T^ (11).

All strains were grown at 28°C in TY medium (12), following which DNA was extracted using Monarch Genomic DNA Purification Kits (New England Biolabs) according to the manufacturer’s instructions. Oxford Nanopore Technologies (ONT) sequencing was performed using Rapid Barcoding Kits (SQK-RBK004, ONT) and R9.4.1 flow cells on a minION device. Basecalling and demultiplexing was performed using Guppy version 6.3.4+cfaa134 (LMG 24692^T^) or 6.4.6+ae70e8f (LMG 27395^T^ and CIP 109850^T^) with the super accuracy mode (ONT). Illumina sequencing was performed at SeqCenter (Pittsburgh, PA, USA) using the Illumina DNA Prep kit, and sequenced on an Illumina NextSeq 2000 to produce 2×151bp reads. Illumina reads were filtered using BBduk version 38.96 (13) and then trimmed using Trimmomatic version 0.39 (14). Sequencing statistics are provided in Table 1.

**Table 1.** Accession numbers, and sequencing, assembly, and annotation statistics.

|  | *Sinorhizobium garamanticum*<br>LMG 24692<sup>T</sup> | *Sinorhizobium numidicum*<br>LMG 27395<sup>T</sup> | *Sinorhizobium numidicum*<br>CIP 109850<sup>T</sup> |
| --- | --- | --- | --- |
| BioProject accession no. | PRJNA935862 | PRJNA935862 | PRJNA935862 |
| BioSample accession no. | SAMN33397752 | SAMN33397753 | SAMN33397754 |
| GenBank accession no. | CP120373 | CP120370 | CP120367 |
|  | CP120374 | CP120371 | CP120368 |
|  | CP120375 | CP120372 | CP120369 |
| ONT reads SRA accession no. | SRR23576515 | SRR23576513 | SRR23576511 |
| Illumina reads SRA accession no. | SRR23576516 | SRR23576514 | SRR23576512 |
| Total ONT read length (nt) | 625,602,711 | 344,577,201 | 1,137,159,815 |
| No. of ONT reads | 69,663 | 83,380 | 315,111 |
| ONT N50 read length (nt) | 15,783 | 7,853 | 6,421 |
| Total Illumina read length (nt) | 3,426,040,094 | 1,987,185,289 | 1,660,861,335 |
| No. of Illumina paired reads | 14,645,955 | 8,521,205 | 7,011,977 |
| Illumina read length (nt) | 2 x 151 | 2 x 151 | 2 x 151 |
| Genome size (bp) | 6,834,248 | 6,722,813 | 6,722,812 |
| No. of protein coding genes | 6,111 | 5,983 | 5,981 |
| G+C content (%) | 61.33 | 60.68 | 60.68 |
| No. of replicons | 3 | 3 | 3 |
| Replicon sizes (bp) | 4,218,344 | 3,768,519 | 3,768,519 |
|  | 1,992,739 | 2,324,848 | 2,324,848 |
|  | 623,165 | 629,446 | 629,445 |

Draft genomes for all strains were generated from the ONT reads using Flye version 2.9-b1779 (15) and contigs less than 2,000 bp removed. The assemblies were then polished using the ONT reads and medaka version 1.7.2 (ONT). Assemblies were further polished using the Illumina reads, first with Polypolish version 0.5.0 (16) and then by POLCA version 4.0.9 (17); in both cases, read mapping was performed using bwa version 0.7.17-r1198-dirty (18). Finally, replicons were reoriented with Circlator version 1.5.5 (19) prior to annotation using PGAP version 2022-10-03.build6384 (20). Assembly and annotation statistics are provided in Table 1.

*S. garamanticum* LMG 24692^T^ and *S. numidicum* LMG 27395^T^ and CIP 109850^T^ each contain multipartite genomes consisting of three circular replicons. As with other *Sinorhizobium* species, the common nodulation and nitrogen fixation genes of both species are found on large symbiotic megaplasmids of ∼623 kb (*S. garamanticum*) and ∼629 bp (*S. numidicum*), respectively. Pairwise average nucleotide identity comparisons against all other *Sinorhizobium* species type strains, as calculated with FastANI version 1.33 (21), confirmed *S. garamanticum* and *S. numidicum* to be distinct species as all values were below 95%. A core-genome maximum likelihood phylogeny was constructed as described previously (Figure 1) (7). The results indicate that *S. garamanticum* forms a clade with *S. terangae, S. mexicanum*, and *S. psoraleae*. On the other hand, *S. numidicum* formed its own lineage, a result that differs from a previous phylogeny created using 16S rRNA gene sequences (11).

**Figure 1.**
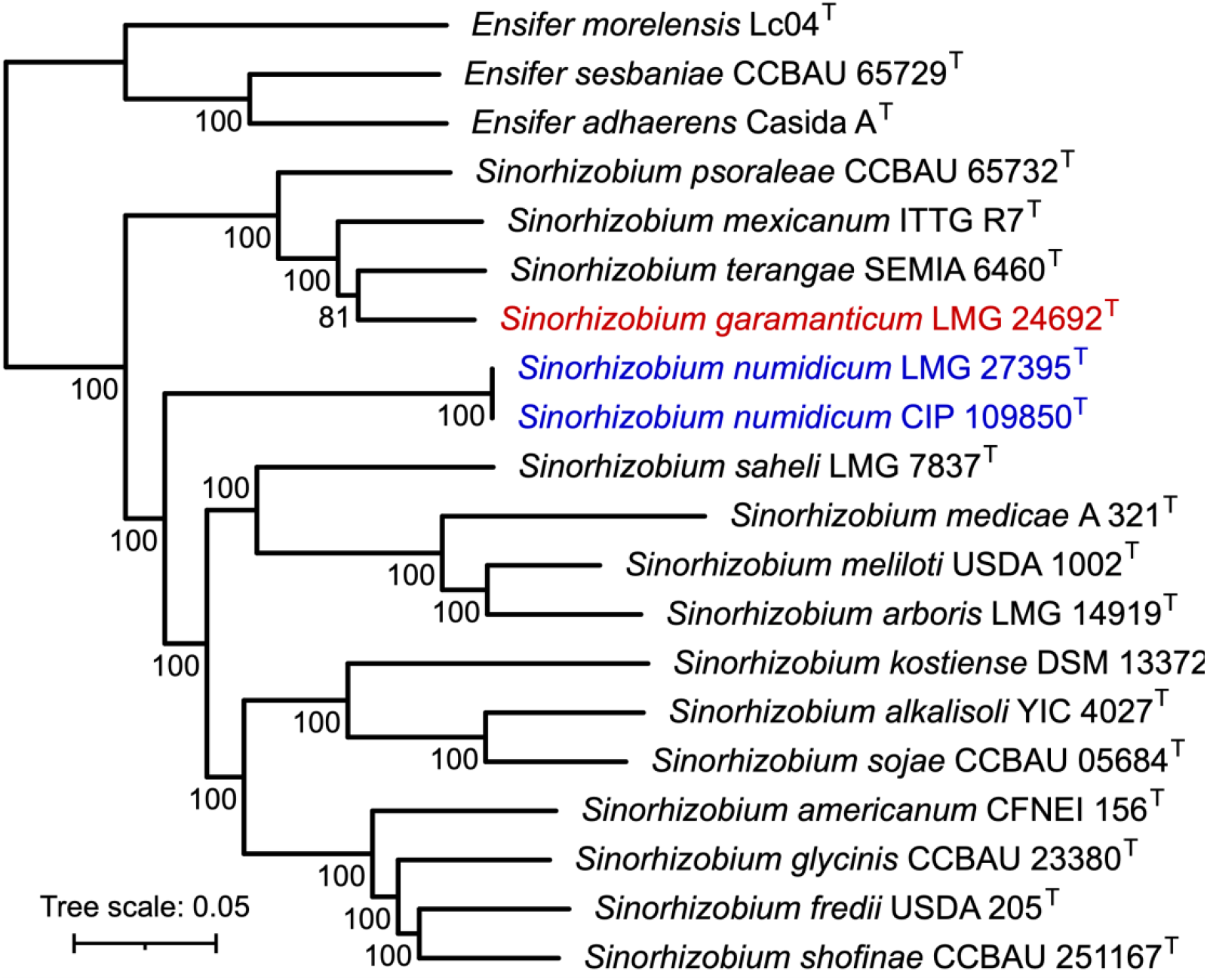
Maximum likelihood phylogeny of the genus *Sinorhizobium*. A maximum likelihood phylogeny of 16 species type strains from the genus *Sinorhizobium* was prepared from a concatenated alignment of 1,509 core genes. *S. garamanticum* is shown in red while *S. numidicum* is shown in blue. Three *Ensifer* spp. (7, 22) were included as an outgroup. To construct the phylogeny, Roary version 1.7.8 (23) was used to identify core genes (80% identity threshold) and to create a concatenated alignment using PRANK version 170427 (24). The concatenated alignment of 1,509 genes was then trimmed with trimAl version 1.4.rev22 (25) and used to construct a maximum likelihood phylogeny using IQ-TREE version 2.2.0 and the GTR+F+I+I+R5 model (26). Values at the nodes represent ultrafast jackknife support values calculated from 1000 replicates and a subsampling proportion of 40%. The scale represents the mean number of nucleotide substitutions per site.

## Data Availability

Annotated genome assemblies were deposited to the NCBI database under BioProject accession <u>PRJNA935862</u>. Raw sequencing reads were deposited to the Short Read Archive and are accessible via the accessions listed in Table 1. Scripts to repeat the genome assembly and annotation, as well as the ANI and phylogenetic analyses, are available through GitHub at github.com/diCenzo-Lab/008_2023_Sinorhizobium_species_type_strains.

## Acknowledgements

Research in the GCD laboratory is supported by the Natural Sciences and Engineering Research Council of Canada through the Discovery Grants program. DES acknowledges “Programa Investigo”, funded by Junta de Castilla y León and “Next Generation EU”. EM acknowledges a European Union’s Horizon 2020 Marie Sklodowska-Curie Actions (Grant Agreement no. 897795).

## References

1. Kuzmanović N, Fagorzi C, Mengoni A, Lassalle F, diCenzo GC. 2022. Taxonomy of Rhizobiaceae revisited: proposal of a new framework for genus delimitation. Int J Syst Evol Microbiol 72:005243.

2. Li Y, Yan J, Yu B, Wang ET, Li X, Yan H, Liu W, Xie Z. 2016. Ensifer alkalisoli sp. nov. isolated from root nodules of Sesbania cannabina grown in saline–alkaline soils. Int J Syst Evol Microbiol 66:5294–5300.

3. Yan H, Yan J, Sui XH, Wang ET, Chen WX, Zhang XX, Chen WF. 2016. Ensifer glycinis sp. nov., a rhizobial species associated with species of the genus Glycine. Int J Syst Evol Microbiol 66:2910–2916.

4. Reeve W, Tian R, Bräu L, Goodwin L, Munk C, Detter C, Tapia R, Han C, Liolios K, Huntemann M, Pati A, Woyke T, Mavrommatis K, Markowitz V, Ivanova N, Kyrpides N, Willems A. 2014. Genome sequence of Ensifer arboris strain LMG 14919T; a microsymbiont of the legume Prosopis chilensis growing in Kosti, Sudan. Stand Genomic Sci 9:473–483.

5. Sugawara M, Epstein B, Badgley BD, Unno T, Xu L, Reese J, Gyaneshwar P, Denny R, Mudge J, Bharti AK, Farmer AD, May GD, Woodward JE, Médigue C, Vallenet D, Lajus A, Rouy Z, Martinez-Vaz B, Tiffin P, Young ND, Sadowsky MJ. 2013. Comparative genomics of the core and accessory genomes of 48 Sinorhizobium strains comprising five genospecies. Genome Biol 14:R17.

6. Lloret L, Ormeño-Orrillo E, Rincón R, Martínez-Romero J, Rogel-Hernández MA, Martínez-Romero E. 2007. Ensifer mexicanus sp. nov. a new species nodulating Acacia angustissima (Mill.) Kuntze in Mexico. Syst Appl Microbiol 30:280–290.

7. Fagorzi C, Ilie A, Decorosi F, Cangioli L, Viti C, Mengoni A, diCenzo GC. 2020. Symbiotic and nonsymbiotic members of the genus Ensifer (syn. Sinorhizobium) are separated into two clades based on comparative genomics and high-throughput phenotyping. Genome Biol Evol 12:2521–2534.

8. Chen WH, Yang SH, Li ZH, Zhang XX, Sui XH, Wang ET, Chen WX, Chen WF. 2017. Ensifer shofinae sp. nov., a novel rhizobial species isolated from root nodules of soybean (Glycine max). Syst Appl Microbiol 40:144–149.

9. Tian CF, Zhou YJ, Zhang YM, Li QQ, Zhang YZ, Li DF, Wang S, Wang J, Gilbert LB, Li YR, Chen WX. 2012. Comparative genomics of rhizobia nodulating soybean suggests extensive recruitment of lineage-specific genes in adaptations. Proc Natl Acad Sci USA 109:8629–8634.

10. Chun J, Oren A, Ventosa A, Christensen H, Arahal DR, da Costa MS, Rooney AP, Yi H, Xu X-W, De Meyer S, Trujillo ME. 2018. Proposed minimal standards for the use of genome data for the taxonomy of prokaryotes. Int J Syst Evol Microbiol 68:461–466.

11. Merabet C, Martens M, Mahdhi M, Zakhia F, Sy A, Le Roux C, Domergue O, Coopman R, Bekki A, Mars M, Willems A, de Lajudie P. 2010. Multilocus sequence analysis of root nodule isolates from Lotus arabicus (Senegal), Lotus creticus, Argyrolobium uniflorum and Medicago sativa (Tunisia) and description of Ensifer numidicus sp. nov. and Ensifer garamanticus sp. nov. Int J Syst Evol Microbiol 60:664–674.

12. diCenzo GC, MacLean AM, Milunovic B, Golding GB, Finan TM. 2014. Examination of prokaryotic multipartite genome evolution through experimental genome reduction. PLOS Genet 10:e1004742.

13. Bushnell B. 2014. BBMap: A Fast, Accurate, Splice-Aware Aligner.

14. Bolger AM, Lohse M, Usadel B. 2014. Trimmomatic: a flexible trimmer for Illumina sequence data. Bioinformatics 30:2114–2120.

15. Kolmogorov M, Yuan J, Lin Y, Pevzner PA. 2019. Assembly of long, error-prone reads using repeat graphs. Nat Biotechnol 37:540–546.

16. Wick RR, Holt KE. 2022. Polypolish: Short-read polishing of long-read bacterial genome assemblies. PLoS Comput Biol 18:e1009802.

17. Zimin AV, Marçais G, Puiu D, Roberts M, Salzberg SL, Yorke JA. 2013. The MaSuRCA genome assembler. Bioinformatics 29:2669–2677.

18. Li H, Durbin R. 2009. Fast and accurate short read alignment with Burrows-Wheeler transform. Bioinformatics 25:1754–1760.

19. Hunt M, Silva ND, Otto TD, Parkhill J, Keane JA, Harris SR. 2015. Circlator: automated circularization of genome assemblies using long sequencing reads. Genome Biol 16:294.

20. Tatusova T, DiCuccio M, Badretdin A, Chetvernin V, Nawrocki EP, Zaslavsky L, Lomsadze A, Pruitt KD, Borodovsky M, Ostell J. 2016. NCBI prokaryotic genome annotation pipeline. Nucleic Acids Res 44:6614–6624.

21. Jain C, Rodriguez-R LM, Phillippy AM, Konstantinidis KT, Aluru S. 2018. High throughput ANI analysis of 90K prokaryotic genomes reveals clear species boundaries. Nat Commun 9:7200.

22. Williams LE, Baltrus DA, O’Donnell SD, Skelly TJ, Martin MO. 2017. Complete genome sequence of the predatory bacterium Ensifer adhaerens Casida A. Genome Announc 5:e01344–17.

23. Page AJ, Cummins CA, Hunt M, Wong VK, Reuter S, Holden MTG, Fookes M, Falush D, Keane JA, Parkhill J. 2015. Roary: rapid large-scale prokaryote pan genome analysis. Bioinformatics 31:3691–3693.

24. Löytynoja A. 2014. Phylogeny-aware alignment with PRANK., p. 155–170. In Russell, D (ed.), Methods in Molecular Biology (Methods and Protocols). Humana Press, Totowa, NJ.

25. Capella-Gutiérrez S, Silla-Martínez JM, Gabaldón T. 2009. trimAl: a tool for automated alignment trimming in large-scale phylogenetic analyses. Bioinformatics 25:1972–1973.

26. Minh BQ, Schmidt HA, Chernomor O, Schrempf D, Woodhams MD, von Haeseler A, Lanfear R. 2020. IQ-TREE 2: new models and efficient methods for phylogenetic inference in the genomic era. Mol Biol Evol 37:1530–1534.

